# NN-RNALoc: neural network-based model for prediction of mRNA sub-cellular localization using distance-based sub-sequence profiles

**DOI:** 10.1101/2021.10.06.463397

**Authors:** Negin Sadat Babaiha, Rosa Aghdam, Changiz Eslahchi

## Abstract

Localization of messenger RNAs (mRNA) as a widely observed phenomenon is considered as an efficient way to target proteins to a specific region of a cell and is also known as a strategy for gene regulation. The importance of correct intracellular RNA placement in the development of embryonic and neural dendrites has long been demonstrated in former studies. Improper localization of RNA in the cell, which has been shown to occur due to a variety of reasons, including mutations in trans-regulatory elements, is also associated with the occurrence of some neuromuscular diseases as well as cancer. We propose NN-RNALoc, a neural network-based model to predict the cellular location of mRNAs. The features extracted from mRNA sequences along with the information gathered from their proteins are fed to this prediction model. We introduce a novel distance-based sub-sequence profile for representation of RNA sequences which is more memory and time efficient and comparying to the k-mer frequencies, can possibly better encode sequences when the distance k increases. The performance of NN-RNALoc on the following benchmark datsets CeFra-seq and RNALocate, is compared to the results achieved by two powerful prediction models that were proposed in former studies named as mRNALoc and RNATracker The results reveal that the employment of protein-protein interaction information, which plays a crucial role in many biological functions, together with the novel distance-based sub-sequence profiles of mRNA sequences, leads to a more accurate prediction model. Besides, NN-RNALoc significantly reduces the required computing time compared to previous studies. Source code and data used in this study are available at: https://github.com/NeginBabaiha/NN-RNALoc

## Introduction

The localization of RNA within the cell has an important role in the regulation of gene expression and has been suggested by numerous studies as a mechanism by which cells are polarized ([1]). Besides, the mRNA localization might be preferable to protein localization as one mRNA molecule can be like a template to code for plenty of proteins and thereby localizing mRNA rather than protein to the site where the protein is needed, is more efficient and time saving. The RNA localization is mainly carried out by a number of regulatory factors such as RNA-binding proteins that can control the distribution of mRNA in each compartment within the cells ([2–5]). To date, the well-known examples of mRNA localization all involve transcripts whose protein products have remarkable roles within specific sub-cellular compartments such as the mRNA encoding the transcriptional repressor ASH1 in budding yeast([6]). Further, in oligodendrocytes, the mRNA encoding myelin basic protein (MBP) is transported into the distal processes where myelination occurs ([7]). Nowadays, studies indicate that the localization of mRNAs to particular sub-cellular compartments may be much more prevalent than previously thought. In a recent study involving high-throughput, high resolution in situ hybridizations of over 3,000 transcripts in Drosophila embryos, 71 were found to be expressed in spatially distinct patterns ([8]). Similarly, in mammalian neurons, it was once thought that only a handful of mRNAs localized at synapses. However, more recent studies indicate that hundreds of mRNAs are present in neuronal processes, where they encode diverse functionalities ([9, 10]). Further, the analysis of RNA localization in migrating fibroblasts ([11]) and Drosophila embryos ([8]) may reveal sub-cellular compartments that had previously been unappreciated, and thus these findings may lead to a more detailed and nuanced understanding of cellular architecture.

The relationship between the location of mRNAs and the proteins that they produce is usually complex and intertwined. Both types of molecules have signals in their sequences which can play a role in transferring them to specific locations within the cell ([12, 13]). As revealed by proteomic analyses, mRNAs to be transported are packaged within complexes containing a large number of associated proteins ([14]). Some of these proteins bind to the mRNA upon transcription or splicing, leading to the future recruitment of the cytoplasmic transport machinery ([15]). Improper localization of RNA in the cell is also associated with the occurrence of some neuromuscular diseases and cancers ([16]). Formerly, a new class of drugs called oligonucleotides has been introduced that target the diseases-causing RNAs rather than proteins ([17–19]). Therefore, improving the understanding of the mRNA localization within the cell is beneficial in the fields such as drug design ([20]). Though mRNA localization has long been studied, only within recent years the computational predictors which are mainly based on machine learning approaches have been introduced. As instance, in a study in 2019, the very first prediction model named as RNATracker was developed that studied mRNA Localization [21]. RNATracker uses convolutional neural networks (CNN) combined with a special form of recurrent neural networks named as long short-term memory, briefly known as LSTM, to make prediction on CeFra-seq and APEX-RIP experimental data. CeFra-seq is a technology used to map human transcript abundance in the nucleus, cytosol, endomembranes and insoluble fraction ([22]). APEX-RIP is an alternate technology that maps the transcriptome of the nucleus, cytoplasm, endoplasmic reticulum (ER) and mitochondria ([23]). Results achieved by RNATracker are also compared to a multi layer perceptron-based predictor named as DNN-5Mer and reveal significant outperformance of RNATracker. DNN-5Mer is based on extraction of k-mer representation profiles from sequences and considers all 1-mers to 5-mers resulting in a 1367-dimensional input vector. The architecture is consisted of two hidden layers of size equal to the input dimension, each followed by ReLU activation and dropout. Surprisingly, in their study it’s clear that RNATracker results gain no improvements when the information of secondary structure of mRNAs are also included in the model ([21]). This may be explained by a number of factors like the imperfect ability to accurately characterize secondary structure using RNAplfold ([24]) and secondly the impact of increased dimention of feature vector when the secondary structure data is incorporated in the model and may easily lead to overfitting. Persumably, more condensed encodings(like k-mer profiles) may prove beneficial. In another recent study a predictive model named as RNA-GPS is introduced to predict localization of the transcripts in APEX-RIP datset only [25]. For each transcript, RNA-GPS computes k-mer frequencies for k between 3 to 5 and assigns a probability for each location using a random forest model. In context of RNA localization, there is a widely-used database named as RNALocate ([26]). RNALocate is a web-accessible database that provides RNA sub-cellular localization resources and documents more than 190,000 RNA-associated sub-cellular localization entries with experimental and predicted evidence. It involves more than 105,000 RNAs with 44 sub-cellular locations in 65 species, mainly including Homo sapiens, Mus musculus, and Saccharomyces cerevisiae. The computational method mRNALoc which is recently investigated in a study ([27]), is mainly trained on a part of RNALocate dataset. mRNALoc is a novel machine-learning based tool that predicts mRNA sub-cellular localization by extracting k-mer profiles from mRNA sequences and applying a support vector machine (SVM) ([26, 27]). As reported, the performance of mRNALoc in two location of cytoplasm and nucleus is better comparing to a recently-introduced computational model named as iLoc-mRNA [27]. iLoc-mRNA which is studied and designed by [28], is based on RNALocate dataset, too, and performs two strategies for feature selection and uses a SVM model for multi-class classification ([28]). mRNALoc has some advantages over RNA-GPS and iLoc-mRNA. First of all data produced by APEX-RIP is roughly noisy and thereby there would be highly beneficial if these later methods assessed their performance on RNALocate datsaet, too. Moreover, the mRNALoc was developed from datasets retrieved from RNALocate which contains manually curated mRNA sub-cellular localization information with experimental evidences. Besides, for cytoplasm and nucleus the performance of mRNALoc was better than iLoc-mRNA but, in endoplasmic reticulum the performance of iLoc-mRNA was better than mRNALoc. It is also noticeable to mention that in iLoc-mRNA prediction were made for one of the following locations namely, cytosol/cytoplasm, ribosome, endoplasmic reticulum, and nucleus/exosome/dendrite/mitochondrion and presumably combining nucleus, exosome, dendrite, and mitochondria as a single location is not appropriate as these are diverse locations which should not be merged in a single sub-cellular class. In this study, we introduce a novel representation from mRNA sequences based on sub-sequences of distance k and we represent that using this encoding along with canonical k-mer frequency profiles, can possibly gather more sequence-based information from mRNAs. We develop a neural network-based model, named as NN-RNALoc that also takes advantage of the protein-protein interactions (PPI) information. Evidently, the PPI information is essential for many cellular activities. Proteins usually function through a number of interactions with other biomolecules and are widely used in biological problems to discover many functions. As an instance, in RNA metabolism and the context of discovering RNA binding proteins (RBPs), a successfull approach predicts RNA binding activity by analyzing protein-protein interactomes without relying on sequence or structure information ([29]). The importance of protein sub-cellular localization is due to the importance of protein’s functions in different cell parts. Moreover, prediction of protein sub-cellular locations helps to identify the potential molecular targets for drugs and has an important role in genome annotation ([30]). All this information and the fact that the proteins with the similar protein-protein interaction patterns tend to be located mostly in the similar sub-cellular location ([30], lead us to take advantage of including this widely-used data in the predictive model.

Besides, it’s also worth noting that to develop a more practical predictor for a biological system, one can also follow “Chou’s 5-steps rule” ([31]). Therefore, in this study, we go through the following five steps to establish the NN-RNALoc predictor ([32, 33]):

1. Select or construct a valid benchmark dataset to train and test the predictor. This step is described in section Data Sources.
2. Represent the mRNAs with an effective formulation and extract k-mer information that can reflect their intrinsic correlation with the target to be predicted. This step is described in Feature Encoding.
3. Introduce and develop the powerful NN-RNALoc algorithm to conduct the prediction. The three main steps of NN-RNALoc and its work-flow are described in NN-RNALoc section.
4. Properly perform cross-validation tests to objectively evaluate the anticipated prediction accuracy. This section is covered in the third step of NN-RNALoc work-flow described also in NN-RNALoc section.
5. In our future work, we will establish a user-friendly web-server for the predictor that is accessible to the public. The more details are stated in the Conclusion section.

As mentioned in [31], the guidelines of Chou’s 5-step rules have the following notable merits:

1. Crystal clear in logic development.
2. Completely transparent in operation.
3. Easily to repeat the reported results by other investigators.
4. With high potential in stimulating other sequence-analyzing methods.
5. Very convenient to be used by the majority of experimental scientist.

NN-RNALoc is a neural network-based predictive model and also consistant with Chou’s 5-steps. It is based on the Cefra-seq and RNALocate resources. The rest of the paper is organized as follows: In the Materials and Methods section, we first declare the datasets and also introduce the features that extracted from mRNA transcripts. Then, we describe the details and steps required to build our machine learning based model. In the Results section, we report the performance of NN-RNALoc on the two mentioned datasets and also compare it with the following two state-of-art methods: mRNALoc and RNATracker. The performance of NN-RNALoc after 10-fold cross-validation is also compared to a neural network-based baseline method named as DNN-5Mer which is formerly executed on Cefra-seq dataset. Finally, in the Discussion section, we extend our experiments to non-human related transcripts and also evaluate the performance of NN-RNALoc using novel distance-based sub-sequence profiles and canonical k-mer information on human and non-human transcripts.

## Materials and methods

Here, we introduce the data sources employed in our research. Then, the details of the information and features are explained and finally the architecture and methodological aspects of NN-RNALoc are described in depth.

### Data Sources

#### mRNA Sequences and Localization Information

Two benchmark datasets are considered. The first dataset is CeFra-seq which is also employed in RNATracker method. As mentioned above, Cefra-seq contains the human transcripts and localization information of mRNAs is presented in the form of normalized gene expression values in each of four sub-cellular locations: cytosol, nuclear, membranes, and insoluble. Hence, for each mRNA, rather than a single cellular location label, we have a vector of length 4 whose elements represent the probability of belonging to each location. The total number of mRNAs is 11,373 and the sequences are downloaded from the Ensemble database ([34]).

In the second dataset, mRNA sequences and sub-cellular localization information are extracted from RNALocate dataset. The same training dataset employed in mRNALoc method is used in this paper, however, NN-RNALoc considers each human and non-human transcripts separately and also for each gene only one isoform is taken into account. In this study only the mRNA sequences that belong to a single location are considered, and it covers five sub-cellular compartments: cytoplasm, endoplasmic reticulum, extracellular region, mitochondrial, and nucleus. The total number of mRNAs here is 11,124 where 5,880 of them are human transcripts and 5,244 of them belong to non-human transcripts. The summary of this dataset is shown in Table 1.

**Table 1.** Total number of mRNAs in each five locations in RNALocate dataset. The first column represents each cellular compartment. The second and third column reveal the number of human and non-human transcripts, respectively.

| Location | Human Species | Non-human species |
| --- | --- | --- |
| Cytoplasm | 3,419 | 1,509 |
| Reticulum Endoplasmic | 1,168 | 6 |
| Extracellular Reigion | 26 | 506 |
| Mitochondria | 2 | 344 |
| Nuclear | 1,265 | 2,869 |
| Total | 5,880 | 5,244 |

#### Protein-Protein Interaction (PPI) Information

The PPI information for human-related mRNAs is extracted from STRING database ([35]). In this database, it usually considers the longest protein-coding isoform among all isoforms of a gene. Hence, one protein is assigned to each mRNA and we keep this information in the form of an adjacency matrix. Each element of this matrix represents the strength of the interaction between two proteins which is assigned by STRING.

### Feature encoding

Two types of features are adopted from the mRNA sequences. The first one is k-mer representation which is one of the most frequently used encodings for nucleotide sequences ([36–38]). The second one is a novel suggested representation of mRNA sequences. The details of these two features are described as follows:

#### k-mer representation

With the explosive growth of biological sequences in the post-genomic era, one of the most important but also most difficult problems in computational biology is how to express a biological sequence with a discrete model or a vector, yet still keep considerable sequence-order information or key pattern characteristics. This is because all the existing machine-learning algorithms can only handle vectors as elaborated in a comprehensive review. One of the ways to extract a uniform length feature vector from these sequences is to count k-mer frequencies ([36, 39]). A k-mer refers to a possible subsequences of length k in a mRNA sequence. As we have 4 neucliotides, the total number of possible k-mers is 4^*k*^.

As an example, for the ACGCCGC sequence, all 5-mer structures are: ACGCC, CGCCG, and GCCGC that are also demonstrated in Figure 1. This k-mer features extracted in this paper can be covered by a very powerful web-server called “Pse-in-One” ([40]).

**Fig 1.**
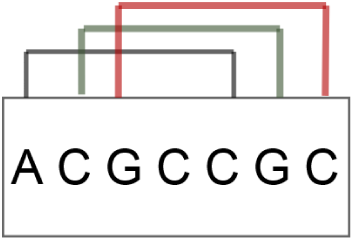
All 5-mer structures contained in ACGCCGC sequence. In this example, we have three 5-mers ACGCC, CGCCG and GCCGC that are shown in three different colors.

For a mRNA sequence S, we first sort all the 5-mer structures lexicographically, then count the occurrence of each one in the main mRNA sequence and divide it by the length of mRNA sequence. Hence, we get the following feature vetor:

*F*_5_(*S*) = [*v*_1_, *v*_2_, …, *v*_*n*_] where each *v*_*i*_ is the frequency of i-th 5-mer, and in this case n is equal to 4^5^ = 1024.

### Distance-based sub-sequence profiles

One of the drawbacks of k-mer representation is that when k increases, the feature vector becomes extremely large and sparse which can be memory-inefficient and can reduce the performance of model. In this work, to overcome this problem and to extract more information with a smaller feature vector for large values of k, we propose a novel distance-based representation. In this representation, the distance between the first and last nucleotide of the subsequence that we count, is set to be k. Then, for each two nucleotides coming with distance k from each other, the frequency of this subsequence is computed. So, for a mRNA sequence S and the distance k, we get the following feature vector of length 16: *D*_*k*_(*S*) = [*w*_1_, *w*_2_, …, *w*_16_], where each *w*_*i*_ is the frequency of the each distance-based sub-sequence. As an example, for k = 2, all subsequences to count are illustrated in bellow:

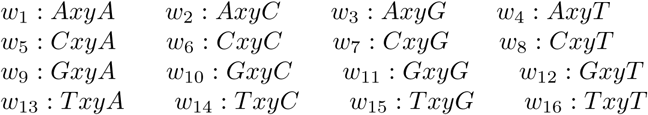

Here, x and y can be substituted with any of four following nucleotides: A, G, C, and T. As an example, let’s consider the S to be the mRNA with sequence ACGCCGC. In this sequence, for *w*_2_ : *AxyC*, only ACGC is contained in the sequence, so frequency of *w*_2_ is 1. For *w*_6_ which is *CxyC*, two subsequences of *CGCC* and *CCGC* are contained in the mRNA sequence and hence the frequency of *w*_6_ is 2. For *w*_11_ which is *GxyG*, only the subsequence *GCCG* is observed and therefore the frequency of *w*_11_ is 1. Other *w*_*i*_s have frequency of zero in this sequence. This example is also illustrated in Figure 2. After experimenting with a variety of ranges for k, we set k to be in range 0 to 8 hence coming up with a feature vector of length 9 ∗ 16 = 144.

**Fig 2.**
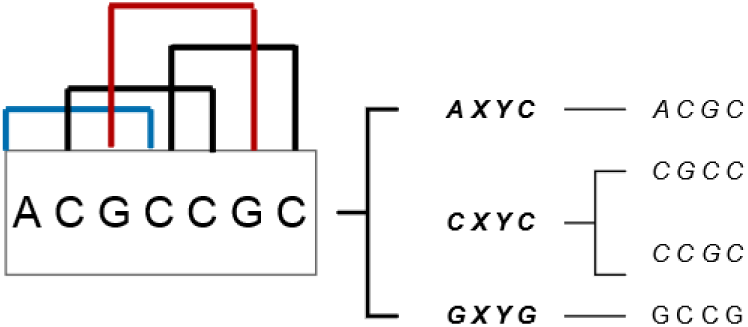
All distance-based sub-structures for k = 2 contained in ACGCCGC sequence. In this example, we have 2 substructure of form: *CxyC*, 1 of form *AxyC* and also 1 of form *GxyG*. Each three distance-based sub-sequence profiles for dsitance k = 2 are shown in three different colors.

### Principle Component Analysis on PPI network

As mentioned above, the PPI information is reflected in the form of the adjacency matrix with the dimention of number of mRNAs ∗ number of mRNAs. Therefore, for the first dataset, the PPI matrix has a dimension of 11, 373 ∗11, 373, and for the human mRNAs in RNALocate dataset, it is a matrix of size of 5880 ∗5880. As performance of machine learning models can degrade when too many features are considered, we first reduce the dimension of this matrix using Principle Component Analysis (PCA) technique ([41]). PCA is among the most widely used methods to reduce the feature space and improve storage space or the computational efficiency of the learning algorithm. It uses singular value decomposition to project the data it to a lower dimensional space and emphasizes variation and brings out strong patterns in a dataset. In this study, after some experiments, the number of principle components is set to 500 and the total explained variance is more than 70 percent of the whole data this way.

### NN-RNALoc

In this section, we express the main steps of NN-RNALoc.

**Step 1:** Aggregation of the following three feature vectors:

1. 5-mer frequencies (a vector of length 1024)
2. Distance-based sub-sequence profiles (a vector of length 144)
3. Reduced PPI matrix using PCA method (a vector of length 500 for each mRNA)

We concatenate the gathered information into a single feature vector of dimension (1024 +144 + 500). This vector is the final vector which is further fed into our neural network model for the prediction task.

**Step 2:** Design of a neural network model

Based on the features obtained, we propose an artificial neural network (ANN) to assign a probability to each of the location classes for each mRNA. A neural network can be expressed as a series of matrix multiplications interleaved with non-linear function. ANN consists of a number of smaller units called neurons which can be repeated in many layers. To prevent the neural network from getting sophisticated and thus becoming more difficult to be trained efficiently, we design a shallow architecture for our model which has one hidden layer and 200 neurons. In the hidden layer, dropout is also used to randomly mask half of the connections during the training of the model to prevent overfitting. We use the rectified linear unit (Relu) activation function in the hidden layer which is defined as below ([42]):

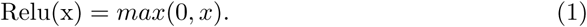

The Softmax function as the non-linear function is applied in the last layer of the model to assign a probability to each location and is formulated as bellow ([43]):

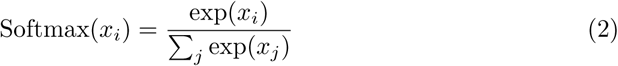

Finaly, we use Kullback-Leiber-Divergence as the loss function. For probability distribution 𝔓 and 𝔔 defined on the same probability space X,

Kullback-Leiber-Divergence is defined as ([44]):

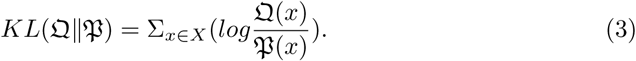

**Step 3:** Training of the prediction model

Hyper-parameters are chosen empirically based on the training dataset. We chose all parameters with the purpose to minimize the loss function. Briefly, we employ the 10-fold cross-validation mode for training the model. Cross-validation is a technique to evaluate predictive models by partitioning the original sample into a training set to train the model, and a test set to evaluate it ([45]). More specifically, In k-fold cross-validation, the data is first partitioned into k equally (or nearly equally) sized segments or folds. Consequently, k iterations of training and validation are performed such that within each iteration a different fold of the data is held-out for validation while the remaining k − 1 folds are used for learning ([45]). We then evaluate the results using a range of values for hidden layers (No hidden layer, 1, 2 and 3), for neurons in each fully connected layer (1000, 700, 500, 200, 100) and for dropout rates (0.1, 0.2, 0.3 and 0.5). The best parameters that are used in this model are shown in Table 2. The number of epochs for training each fold is 300 and a validation set consisting of 10 percent of the training data is also applied to monitor the loss function in the training process and detect overfitting. NN-RNALoc is implemented using the Keras Library ([46]). Adam optimizer with Nesterov momentum is also employed for training the model ([47]). The overall workflow of NN-RNALoc is shown in Figure 3.

**Table 2.** Selected hyperparameters for NN-RNALoc.

| Parameter | Value |
| --- | --- |
| Epochs | 300 |
| Batch size | 512 |
| Hidden layer | 1 |
| Fully connected neurons | 200 |
| Dropout rate | 0.2 |

**Fig 3.**
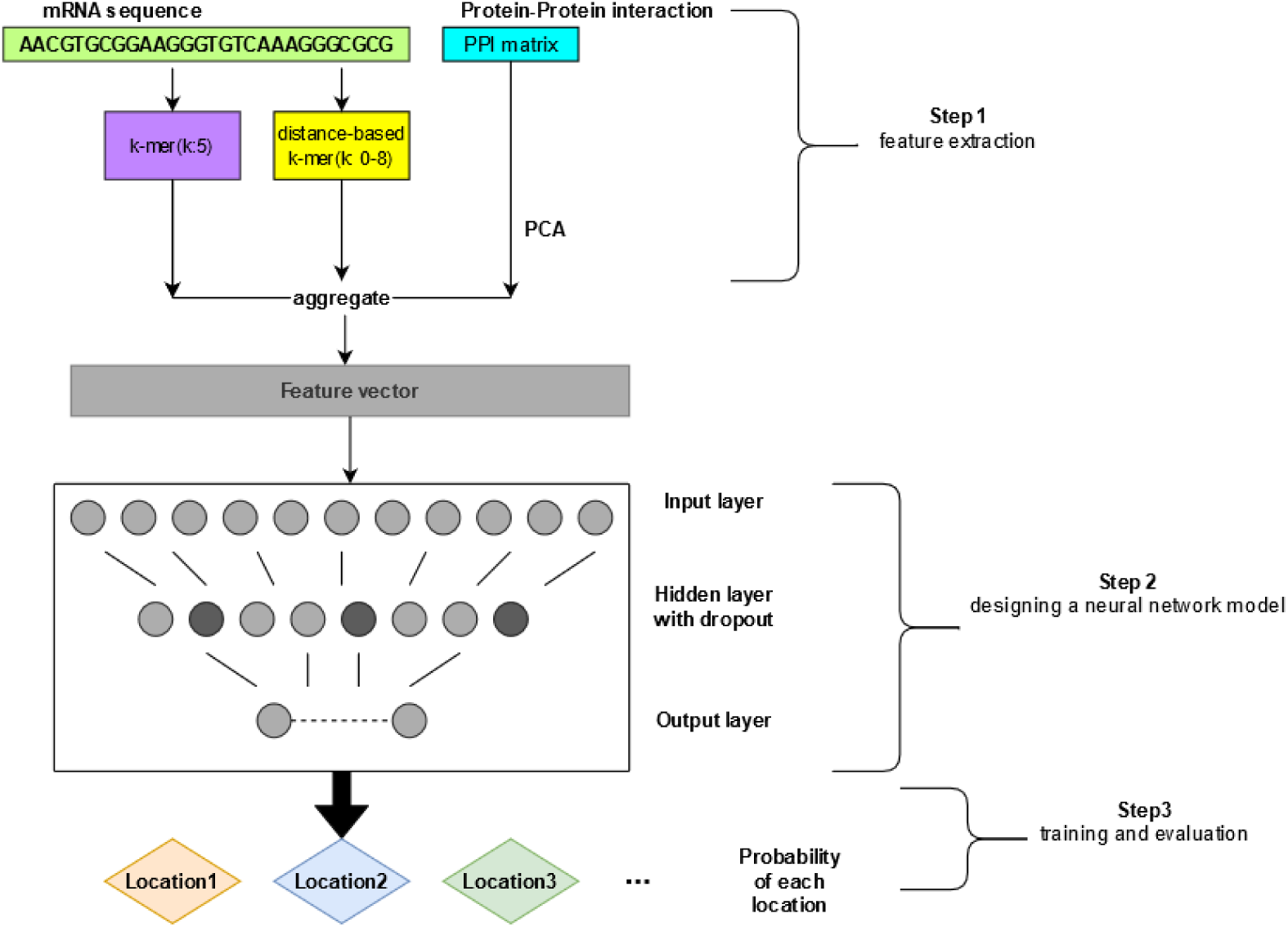
The overflow of NN-RNALoc method. In Step 1, we first aggregate all information gathered from both sequence-based features as well as PPI matrix. In Step 2, we design a neural network model with the mentioned architecture. In Step 3, the model is trained and evaluated using 10-fold cross-validation. For the CeFra-seq dataset, we report a vector of length 4 for each mRNA and for the RNALocate dataset, we report the location with maximum probability to be the predicted location of the mRNA.

## Results

### Time complexity

Many of the methodologies like mRNALoc, are available as a server tool and hence make it impossible to do time complexity related comparisons. Among them, however, RNAtracker’s source code and implementation are available and NN-RNALoc outperforms considerably the RNAtracker method in terms of training time. For NN-RNALoc this time is computed as about 3 hours on Linux Ubuntu machine, with 15 CPUs(Intel Xeon(R) 2.00 GHz) on CeFra-seq dataset and comparing to RNATracker, is noticeably reduced(RNAtracker requires 7 days for training in full length mode and 8 hours in fixed length mode on GTX1080Ti graphic card). On the RNALocate dataset, NN-RNALoc has training time of 2 hours which is again remarkably decreased comparing to RNATracker which needs the computation time of 6 hours. The computation time of DNN-5Mer is roughly the same as NN-RNALoc.

### Evaluation criteria

Pearson correlation is computed to compare the predicted values with experimental ones. Pearson correlation is a measure to assess the linear correlation between variables. It has a value in range (−1, +1). A value of +1 is total positive linear correlation, 0 is no linear correlation, and −1 is total negative linear correlation. To better evaluate the model’s performance, we also consider the Spearman correlation between the predicted and experimental values to capture the order of locations to which a mRNA belongs. For the second dataset, as one location is dedicated to each mRNA, we report the location with the maximum probability to be the final predicted location of mRNA. In order to compare the performance of NN-RNALoc method to other methods for second dataset, True Positive (TP), True Negative (TN), False Positive (FP), False Negative (FN), Precision, Recall, F-score, Accuracy (ACC) and Matthews Correlation Coefficient (MCC) are calculated as follows:

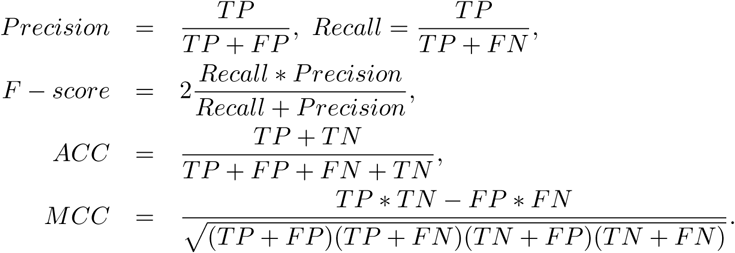

The criteria mentioned above are among the most commonly used metrics in classification problems that can define how good a model has performed. As we know, assessment of some statistical measures alone (like Accuracy) can show overoptimistic inflated results, especially on imbalanced datasets. Therefore, in our results we also evaluate improvements to MCC which is a more reliable statistical rate and produces a high score only if the prediction obtained good results in all categories.

### Assessment and Comparison

The average results of 30 times 10-fold cross-validation mode of training on the first dataset are shown in Table 3. In the CeFra-seq dataset, NN-RNALoc obtains the Pearson correlations of 0.69, 0.65, 0.54 and 0.55 in cytosol, insoluble, membrane and nuclear, respectively. These correlations are higher in all locations comparing to RNATracker fixed length mode method. For total 2893 number of mRNAs in this dataset, the Spearman correlation between the real and predicted vectors is computed as 1 which is again slightly better comparing to RNATracker(2849). As Table 3 demonstrates, NN-RNALoc achieves about 16% better overall Pearson correlation comparing to RNATracker fixed length mode and about 1% better overall Pearson correlation compared to RNATracker full-length mode. As mRNALoc is available as a standalone tool which is trained based on five different locations than Cefra-seq, it was not feasible to take it into the comparison with NN-RNALoc on this dataset. We also compare the performance of NN-RNALoc with DNN-5Mer. As mentioned in the former sections, DNN-5Mer extracts k-mer features and has only two hidden layers with the same number of neurons as the input vector. It also uses Relu activation function in the hidden layer. Though both NN-RNALoc and DNN-5Mer have a simple architecture, however, the performance of DNN-5Mer is noticeably weak and it achieves the Pearson correlation of 0.63 in cytosol, 0.55 in insoluble, 0.42 in membrane and 0.48 in nuclear and in overal, NN-RNALoc achieves about 34 percent higher Pearson correlation comparing to DNN-5Mer. The results can reveal the benefits of employing informative sequence-based information as well as PPI data in the prediction model for mRNA localization while utilizing shallow neural network architecture.

**Table 3.**
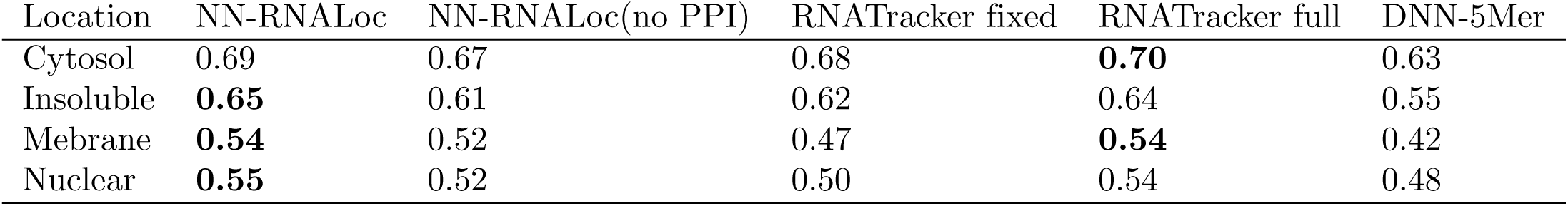
Average Pearson correlations of 30 times 10-fold cross-validation in each location using three methods: NN-RNALoc(with and without employing PPI), RNATracker(fixed and full length mode) and DNN-5Mer on CefraSeq daatset.

The performance of NN-RNALoc on the second dataset is also shown in the Table 4. As illustrated in this table, in cytosol NN-RNALoc has Precision, Recall and F-score of 74%, 72%, and 74%, respectively. In Endoplasmic reticulum, the Precision is 56% and Recall is 48% while F-score is 52%. In Extracellular region and mitochondria, due to few training samples(only 26 and 2, respectively) belonging to these location, Recall and F-score are all near to zero and the Precision in Extracellular region is 91%. In nucleus, the Precision of prediction is 52%, while Recall and F-score are 70% and 60%, respectively. In three following locations cytosol, ER and EX, NN-RNALoc has improved the average of F-score in all locations by 4% comparing to mRNALoc, and by 12% comparing to RNATracker. In nucleus, however, the average of F-score obtained for NN-RNALoc is pretty much the same as mRNALoc. The overall Accuracy of prediction using NN-RNALoc is 65%, which is 4% better comparing to mRNALoc and 2% higher when compared to RNATracker(fixed length mode). NN-RNALoc achieves also MCC of 0.40, which is higher comparing to both RNATracker and mRNALoc methods(that achieve MCC of 0.34 and 0.37, respectively.)

**Table 4.** Results of 30 10-fold cross-validation of NN-RNALoc (with and without employing PPI) compared with RNATracker(fixed length mode) and mRNALoc on the human part of RNALocate dataset. The names of compartments are abbreviated as Cyt : cytosol, ER: Endoplasmic reticulum, EX : Extracellular region, Mit :Mitochondria, Nuc: Nucleus.

|  | NN-RNALoc |  |  |  |  | NN-RNALoc(noPPI) |  |  |  |  | RNATracker |  |  |  |  | mRNALoc |  |  |  |  |
| --- | --- | --- | --- | --- | --- | --- | --- | --- | --- | --- | --- | --- | --- | --- | --- | --- | --- | --- | --- | --- |
| Criteria | Cyt | ER | EX | Mit | Nuc | Cyt | ER | EX | Mit | Nuc | Cyt | ER | EX | Mit | Nuc | Cyt | ER | EX | Mit | Nuc |
| Precision | **0.74** | **0.56** | **0.91** | 0.00 | 0.52 | 0.65 | 0.40 | 0.00 | 0.00 | **0.61** | 0.66 | 0.52 | 0.00 | 0.00 | 0.49 | 0.74 | 0.46 | 0.00 | 0.00 | 0.52 |
| Recall | 0.72 | **0.48** | **0.04** | 0.00 | 0.70 | **0.84** | 0.14 | 0.00 | 0.00 | 0.51 | 0.59 | 0.06 | 0.00 | 0.00 | 0.54 | 0.61 | 0.50 | 0.00 | 0.00 | **0.72** |
| F-score | **0.74** | **0.52** | **0.07** | 0.00 | 0.60 | 0.73 | 0.21 | 0.00 | 0.00 | 0.56 | 0.73 | 0.11 | 0.00 | 0.00 | 0.53 | 0.67 | 0.48 | 0.00 | 0.00 | **0.61** |
| Average Accuracy | **0.65** |  |  |  |  | 0.63 |  |  |  |  | 0.63 |  |  |  |  | 0.61 |  |  |  |  |
| Average MCC | **0.40** |  |  |  |  | 0.30 |  |  |  |  | 0.34 |  |  |  |  | 0.37 |  |  |  |  |

## Discussion

In order to assess the impact of including the PPI information in our model, we perform the following analysis on both Cefra-Seq and RNALocate datasets. Here, we employ both 5-mer and distance-based sub-sequence information gathered from mRNA sequences and then, compare the results with the case when PPI information is also used in the model. As Table 3 demonstrates, on Cefra-se dataset when NN-RNALoc employs the reduced PPI matrix in the model, it performs significantly better and achieves 10% higher Pearson correlation in average for all locations. However, it is also important to notice without inclusion of PPI, the performance of NN-RNALoc is still better in cytosol and insoluble comparying to RNATracker(fixed length) and it achieves almost the same Pearson correlation. Besides, using this setting, in membrane and nucleus it achieves 5 and 2 percent higher correlation, individually. This makes it clear that by using only the sequence-based information of mRNAs, NN-RNALoc still outperforms the former method RNATracker and achieves higher overall Pearson correlation. We do the same analysis on the human-related transcripts of RNALocate dataset and consider only sequence-based information in the model. As Table 4 demonstrates, in this case the average of Precision, Recall, F-score, Accuracy and MCC in all locations are computed as 0.33, 0.29 and 0.30, 0.63 and 0.30, respectively. These results represent that in the second dataset, too, when NN-RNALoc employs PPI information, the performance gets clearly better and it achieves average of Precision, Recall, F-score, Accuracy and MCC of 0.55, 0.38, 0.37, 0.65 and 0.40, respectively.

Although by employing sequence-based features alone, NN-RNALoc still has better performance compared to mRNALoc and RNATracker, the improvement in the performance of NN-RNALoc when it considers the PPI information, may probably highlight the importance of this PPI information and reveal its potential role cellular localization of mRNAs.

At last, we also consider the non-human transcripts of RNALocate dataset. In this case, due to a variety of species whose trasncripts are covered in the dataset, it is not applicable to utalize the PPI information. Therefore, we employ only 5-mer and distance-based sub-sequence profiles of mRNA sequences in the model. The performance of NN-RNALoc on non-human species is described in Table 5 and is compared to mRNALoc and RNATracker. The details and distributions of non-human transcripts of RNALocate dataset are formerly declared in Table 1. The average of Precision, Recall, F-score, ACC and MCC in all locations for NN-RNALoc in this case are 3%, 6%, 6%, 4% and 8% higher comparing to RNATracker, respectively. Comparing to mRNALoc method, too, the average of Precision, ACC and MCC for NN-RNALoc are 6%, 9% and 12% better, separately.

**Table 5.**
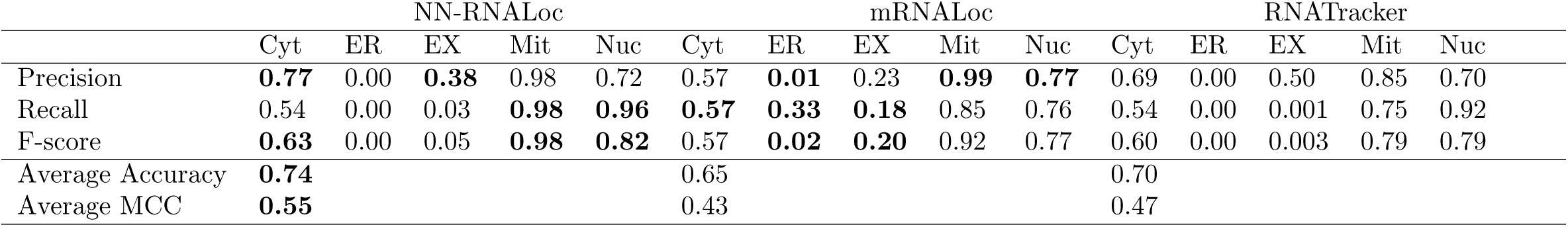
Performance of NN-RNALoc, RNATracker(fixed length mode) and mRNALoc on non-human mRNAs of RNALocate dataset.

|  | NN-RNALoc |  |  |  |  | mRNALoc |  |  |  |  | RNATracker |  |  |  |  |
| --- | --- | --- | --- | --- | --- | --- | --- | --- | --- | --- | --- | --- | --- | --- | --- |
|  | Cyt | ER | EX | Mit | Nuc | Cyt | ER | EX | Mit | Nuc | Cyt | ER | EX | Mit | Nuc |
| Precision | **0.77** | 0.00 | **0.38** | 0.98 | 0.72 | 0.57 | **0.01** | 0.23 | **0.99** | **0.77** | 0.69 | 0.00 | 0.50 | 0.85 | 0.70 |
| Recall | 0.54 | 0.00 | 0.03 | **0.98** | **0.96** | **0.57** | **0.33** | **0.18** | 0.85 | 0.76 | 0.54 | 0.00 | 0.001 | 0.75 | 0.92 |
| F-score | **0.63** | 0.00 | 0.05 | **0.98** | **0.82** | 0.57 | **0.02** | **0.20** | 0.92 | 0.77 | 0.60 | 0.00 | 0.003 | 0.79 | 0.79 |
| Average Accuracy | **0.74** |  |  |  |  | 0.65 |  |  |  |  | 0.70 |  |  |  |  |
| Average MCC | **0.55** |  |  |  |  | 0.43 |  |  |  |  | 0.47 |  |  |  |  |

Eventually, as illustrated by the results, NN-RNALoc has been capable to outperform both methods in most of the criteria in these two datasets. This is probably determinant of the role and importance of explained sequence-based features as well as PPI information in mRNA sub-cellular localization.

## Conclusion

NN-RNALoc stands among the few proposed methods that have applied the machine learning based approaches to study the cellular localization of mRNAs. Facing the explosive growth of biological sequences discovered in the post-genomic age, to timely use them for variety of problems in bioinformatics like RNA and protein localization or drug development, a lot of important sequence-based information, such as PTM (posttranslational modification) sites in proteins, have been successfully predicted ([48]). The rapid development in sequential bioinformatics and structural bioinformatics and introduction of computational methodologies for this means have driven this research area undergoing an unprecedented revolution. In view of this, the computational (or in silico) methods were also utilized in the current study. Localization of mRNA molecules within the cytoplasm provides a basis for cell polarization, thus underlying developmental processes such as asymmetric cell division, cell migration, neuronal maturation and embryonic patterning ([49]). A huge advantage of mRNA targeting is that it allows regulation of gene expression in both space and time and therefore, RNA localziation would have significant advantages for understanding cellular functions ([49]). NN-RNALoc is neural network-based tool which is introduced in this work for prediction of mRNA sub-cellular localization and aims to profit from the interaction information of the proteins coded by mRNA transcripts. We have also suggested an alternate distance-based subsequence profiles for representation of mRNA sequences. This novel encoding which is more condensed and less prone to add redundancy to the data was introduced to address the memory and time related problems appeared in k-mer representation when k increases. The results demonstrate that by employement of the distance-based sub-sequence profiles along with k-mer frequencies and with inclusion of PPI matrix data, NN-RNALoc which has simple and transparent neural network architecture, outperforms two formerly introduced and powerful methods. This simplicity also reduces the required computational time of the training models to a remarkable extent. It is also important to notice that using a dimensional reduction technique such as PCA applied on the PPI data which is a high-dimensional matrix, proves to be highly beneficial. In the future, other dimensionality reduction techniques such as autoencoders or more PPI specific compressing approaches can also be employed. Moreover, it’s precious to notice that inclusion of some other important but rarely-used information such as the knowledge of protein 3D (three-dimensional) structures or their complexes with ligands which is vitally important in many studies in computational biology like drug design ([52]), can be further evaluated in the future studies. Though there are still some difficulties for determining these structures, some tools are introduced such that NMR which is indeed a very powerful tool in determining the 3D structures of membrane proteins([54–58]), albeit it is time-consuming and costly, as well. To acquire the structural information in a timely manner, a series of 3D protein structures have been developed by means of structural bioinformatics tools. For this means, in the future versions of NN-RNALoc, inclusion of structural information of proteins could also be assessed. Besides, as shown in a series of recent publications ([59, 60]) in demonstrating new findings or approaches, user-friendly and publicly accessible web-servers will significantly enhance their impacts ([31, 61]), and so we shall and will definitely make efforts in our future work to provide a web-server to display the findings that can be manipulated by users according to their needs.

## Acknowledgments

Changiz Eslahchi and others would like to thank the School of Biological Sciences, Institute for Research in Fundamental Sciences (IPM) and Computing Center of IPM in performing a parallel computing is gratefully acknowledged.

## Notes

### Competing Interest Statement

The authors have declared no competing interest.

